# A deep-learning based analysis framework for ultra-high throughput screening time-series data

**DOI:** 10.1101/2024.08.22.609110

**Authors:** Patrick Balzerowski, Lukas Hebig, Francisco de Abreu e Lima, Erica Manesso, Thomas Müller, Holger Diedam, David Gnutt

## Abstract

Analysis of ultra-high-throughput screening data sets is a highly critical step in drug discovery campaigns. Due to various environmental and experimental error sources fast and reliable identification of possible candidate compounds is challenging. In this work, we introduce a novel deep-learning based analysis framework to analyze uHTS time-series data sets. Our framework is based on two independent deep-learning models. A deep-learning regression model reduces temporal and spatial signal variation across multitier plates caused by systematic and random errors and a separate variational autoencoder model is used for dimensionality reduction. In contrast to classical evaluation methods our approach is capable to derive lower dimensional representations of time-series signals without a-priori knowledge of the data generating mechanism. We tested our analysis framework on an experimental uHTS data set and identified two distinct classes of substances in the screened library which could be attributed to two biological modes of action. Selected substances belonging to both modes of action were successfully validated in a secondary screening experiment.

## Introduction

Ultra-high-throughput screening (uHTS) is an integral part of drug discovery campaigns. This interdisciplinary technique combines automation, assay-miniaturization and data processing to screen vast chemical libraries and quantify their effect on biological targets^1^. Modern uHTS campaigns use libraries consisting of multiple million compounds^1–4^. The main goal of screening scientists is to identify possible candidate compounds/compound classes rapidly and accurately^1,4,5^.

Most uHTS experiments are either based on biochemical or cell-based assays^6^. Biochemical assays offer a direct way to interrogate a biological target of interest. Compared to cell-based assays miniaturization of biochemical assays is easier accomplished and experiments exhibit less variability due to the homogenous nature of reactions^7^. However, biochemical assays heavily rely on the purification of the biological targets of interest^1^. Besides this, active compounds identified via biochemical assays can lack activity in vivo, due to various requirements enforced by the natural cellular environment of the target^8^. In contrast to this, cell-based assays provide the possibility to test substances directly in the cellular environment enabling the investigation of whole pathways^9^. This potentially allows identification of distinct modes of action of potential drug candidates at the earliest stage of the drug discovery pipeline. Due to this more complex testing environment, experimental responses encountered in cell-based assays (typically transient fluorescence signals) can exhibit a more complex nature compared to mostly binary-type responses in biochemical assays^10^. This potential increase in information content comes at the price of much higher experimental variability, which is mainly caused by an increase in biological variability.

Although signals obtained by cell-based assays potentially contain valuable information related to the biological response or compound action, activity of substances is commonly measured via one- or two-dimensional estimators derived from transient response data^5^. Examples of such estimators are area under the curve (AUC) or integration/summation of subsections of transient signals at specific points in time (point-in-time analysis). In subsequent analysis steps preprocessing of these lower dimensional estimators is performed. Preprocessing aims to correct for known systematic experimental effects and to normalize the data into percentage activity values often using control wells as an internal reference. At the final analysis stage, experimentalists determine suitable decision boundaries based on these calculated estimators, which basically constitute a lower dimensional representation of the full screening data set, to identify primary hits (see schema in Figure 1).

**Figure 1.**
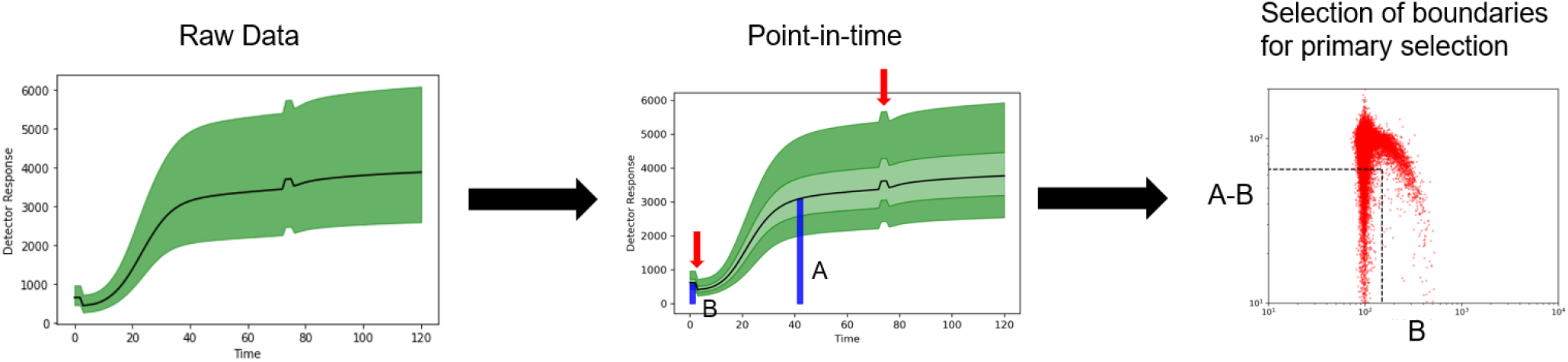
Typical steps in uHTS data analysis.

In the past, many approaches to detect and remove systematic experimental effects from uHTS data sets have been proposed^3,5,11–17^. All of these methods are directly applied to the low-dimensional signal estimators and use statistical methods to remove spatial bias and reduce plate-to-plate drifts in the uHTS data set.

In this article, we want to add a new perspective to the analysis of uHTS data and propose a novel analysis framework based on deep-learning models, which makes use of the entire transient signal as an input. Our framework consists of two individual neural network models. A preprocessing model reduces temporal (plate-to-plate) as well as spatial (across microtiter plates) variation caused by systematic and random errors and a separate representation learning model is used to derive an unbiased dimensionality reduction of the data set without a-priori knowledge of the data generating mechanisms.

## Methods

### Datasets

All plots presented in Figures 3 to 13 are based on an experimental uHTS data. The data is taken from a uHTS screening campaign based on a cell-based calcium flux assay. This assay was optimized to find inhibitors for a biological target. The experimental data set in total contains 2862 microtiter plates of 1536 well format, thus resulting in a total number of 4396032 individual measurements. Each measurement corresponds to a transient fluorescent read-out signal and consists of 121 data points. uHTS experiments were performed in 13 successive batches with an average of 220 plates per batch. In total 549504 neutral control measurements and 3663360 substance measurements have been performed.

To demonstrate the application of our analysis framework on more complex transient signals we generated an additional artificial toy data set. Figures 14 and 15 are based on this artificial data set. In total five batches of 200 plates in 1536 plate format were generated. The toy data set consists of five distinct signal classes which resemble typical shapes encountered in biological experiments.

### Machine learning based data processing

Our deep-learning based analysis framework consists of three steps: preprocessing, dimensionality reduction and scoring. In the following sections each of these steps will be described. All algorithms were implemented in Python using tensorflow to create/train deep-learning models and scikit-learn for our scoring methods based on gaussian mixture models^18,19^.

#### Preprocessing via deep-learning model

The main goal of preprocessing uHTS data sets is to decrease plate to plate variability and to establish reliable assay background levels^16^. Experimentally, within-plate control measurements are included during screens for this purpose^20^. Common sources of signal variance include biological variation and effects due to differences in evaporation rate across individual plates (edge-effect)^21^. In addition to this, systematic errors can be introduced by procedural effects e.g. errors in pipetting processes during liquid handling^16^. Depending on the number of substances which need to be tested during a campaign (typically up to several million compounds), uHTS experiments can last multiple weeks. Thus, besides basic signal variation also intra-batch drifts and batch-to-batch offsets in signal levels for consecutively measured plates are commonly encountered^22,23^.

In classical HTS data processing normalization is typically not performed at raw data level but only after point-in-time analysis^5,16^. In contrast to this, our preprocessing step is directly applied to the log-transformed raw transient signals. Here, we employ a deep-learning regression model for preprocessing. The basic architecture of the model which is a standard feedforward neural network with fully connected layers, is summarized in Table 1.

**Table 1.** Preprocessing model architecture and parameterization.

| Layer | Size | Activation | Dropout |
| --- | --- | --- | --- |
| Input Meta-Data | 4 | Linear | NA |
| Dense | 256 | ReLU | 0.1 |
| Dense | 256 | ReLU | 0.1 |
| Dense | 256 | ReLU | 0.1 |
| Dense | 1 | Linear | NA |
| Input Signal | 121 | Linear | NA |
| Output | 121 | Linear | NA |

Our model is based on the following reasoning: In uHTS experiments most tested substances will be inactive and thus produce an experimental response similar to a neutral control signal. Therefore, we argue that global signal variation across the entire assay data set can be reduced by performing a simple regression of each individual signal towards the global median of neutral control signals. Furthermore, main sources of variation can originate on the one hand from spatial effects (position of a well on an individual plate) and on the other hand be linked to temporal effects (batch-to-batch and intra-batch). Accordingly, our model is trained to infer an additive factor which reduces the MSE with respect to the global median of neutral control signals based on two spatial and two temporal input parameters: spatial input parameters correspond to row/column indices of an individual well on a plate and temporal parameters are given by an intra-batch and batch index. Since our model acts on the log-transformed raw signals, the additive factor inferred by our model acts as a multiplicative factor on the raw data level.

#### Dimensionality reduction by variational autoencoder

As stated above in an uHTS experiment most tested substances are expected to behave similar to neutral control measurements and true primary hits will be rare events^3^. Furthermore, as the assay optimization is performed based on a specific effect on a biological target, hits of various strength intuitively shall be continuously located along a common direction in the lower dimensional encoding. In this work, we employ a modified variational autoencoder model to produce such a lower dimensional encoding of the uHTS data set.

In general, a standard autoencoder is a deep-learning model which consists of two sub-networks, an encoder and a decoder part. The whole autoencoder network is trained to copy its input to its outputs with minimal error^24^. The encoder network maps the input data to a fixed size lower dimensional representation (latent encoding). These latent encodings serve as inputs for the decoder network which aims to map them back to the original input. By constraining the network architecture an information bottleneck is introduced between the encoder and decoder. This forces the encoder to find the salient underlining structure in the data and compress the input data in an efficient way to enable the decoder network to reconstruct the input from the compressed latent encodings. In variational autoencoders encodings generated by the encoder network are not directly used as inputs for the decoder network. Instead, the encoder maps the input to a mean encoding *µ* and a standard deviation encoding *σ* of fixed dimension. The actual encoding provided for the decoder network is then sampled randomly from a gaussian distribution with mean *µ* and standard deviation *σ*. This enables variational autoencoders to learn smooth continuous latent encodings for the input data set.

The basic architecture of the variational autoencoder used in this article is summarized in Figure 2. We basically employ a symmetrical encoder-decoder architecture. The encoder consists of three fully connected layers with 16, 8 and 4 neurons in each layer. The latent encoding is two-dimensional and a gaussian prior is used. The standard loss function for a variational autoencoder contains one part measuring reconstruction error (here MSE between input and reconstructions) and the Kullback-Leibler divergence which measures the distance between the resulting latent distribution and a gaussian prior. For our model we added a third term to the loss function that minimizes the length of mean encoding vectors for control samples. Thereby, we force the encoding to include a point of reference (the origin of the encoding space) given by known control samples which is helpful for defining a scoring system based on the similarity of an individual measurement with respect to the control. A detailed explanation of the loss function used in our variational autoencoder model can be found in the supplementary materials.

**Figure 2.**
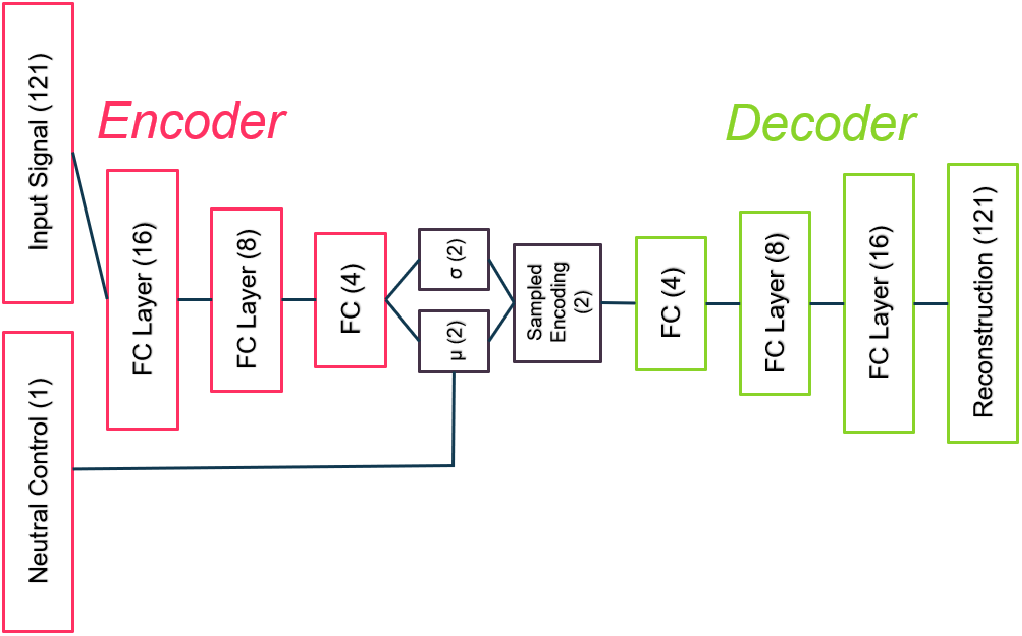
Network architecture of variational autoencoder used for dimensionality reduction.

#### Scoring

Based on the two-dimensional encoding of the data set we employ an anomaly detection approach to quantify the similarity of substance measurements with respect to neutral control measurements. As mentioned in the previous section due to an additional constraint term in the loss function of the autoencoder known neutral control samples are located close to the origin of the encoding space. Our scoring approach builds on this property of the encoding space and consists of two steps. First, using a gaussian mixture model we derive a density estimation for known neutral control measurements. Selection of the number of gaussians needed for this fit is performed by employing an elbow method based on the Bayesian information criterion. Secondly, this fitted gaussian mixture is used to calculate a contrast score for all substance measurements. Basically, instances of substance measurements located at high-density regions of the neutral control distribution receive low contrast scores and instances at low-density regions receive high contrast scores. A detailed explanation of the calculation of contrast scores can be found in the supplementary materials.

Typical uHTS experiments last multiple days/weeks and are carried out batch-wise with on average around 200 to 250 plates per batch. We decided to perform calculation of contrast scores on a batch-to-batch basis in order to establish a frame of reference for each consecutive batch of measured plates. Therefore, contrast scores specifically contain information about the similarity of substance measurements with respect to the distribution of neutral control measurements within a batch. Thereby, we augment the information content beyond the lower dimensional encoding and enable selection of interesting experimental observations not only based on location within the encoding space but also based on contrast scores.

## Results

### Preprocessing model

As stated in the methods section main sources of variation can be divided in two distinct domains. One source is a variation of measurements across individual plates due to differences in e.g. evaporation rate, cell seeding or various other liquid handling effects (spatial plate effects). The second major source of variation is linked to effects which affect the whole experimental setup and stem from the successive measurement of plates. These effects manifest in measurement drifts for successively measured plates or jumps in signal level between batches (temporal effects).

To quantify and visualize the impact of our preprocessing model we consider two metrics: basal-signal-average (*BSA*) and plate-deviation (*PD*). BSA is defined as average of the initial 5 seconds of an individual signal. Based on this PD can be defined in the following way:

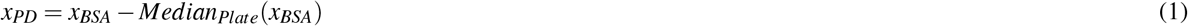

Figure 3 displays plate medians of *BSA* for all successively measured plates of our uHTS experiment. As can be seen from the red curve in Figure 3 most apparent temporal features in the raw data are intra-batch drifts and abrupt jumps of *BSA* level on a batch-to-batch basis. After preprocessing (blue curve) intra-batch drifts as well as inter-batch jumps are significantly reduced. To highlight the effect of our preprocessing model on spatial variations across individual plates consider Figure 4 to 6 which depict the distribution of *PD* values across individual plates. In case of the raw plate-deviation values an accumulation of strongly deviating wells at the edges of the plates is observed. This well-known artifact, commonly referred to as “edge effect”, is mainly caused by increased evaporation rates in the outer wells of a plate. A comparison of the raw data to the preprocessed data clearly shows a reduction of this edge effect. In general, as indicated by the histogram of plate-deviation values, the preprocessing model leads to a narrowing of the distribution and thus increases the homogeneity of signal levels across individual plates.

**Figure 3.**
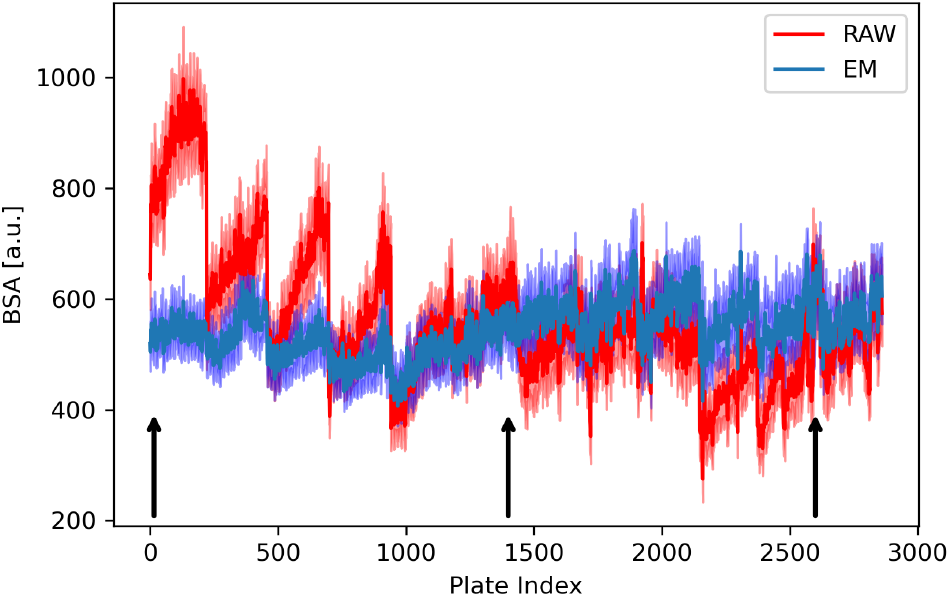
Median of basal-signal-average (BSA) for all successively measured microtiter plates in the uHTS experiment. Shaded region corresponds to the inter-quartile-range of BSA per plate.

**Figure 4.**
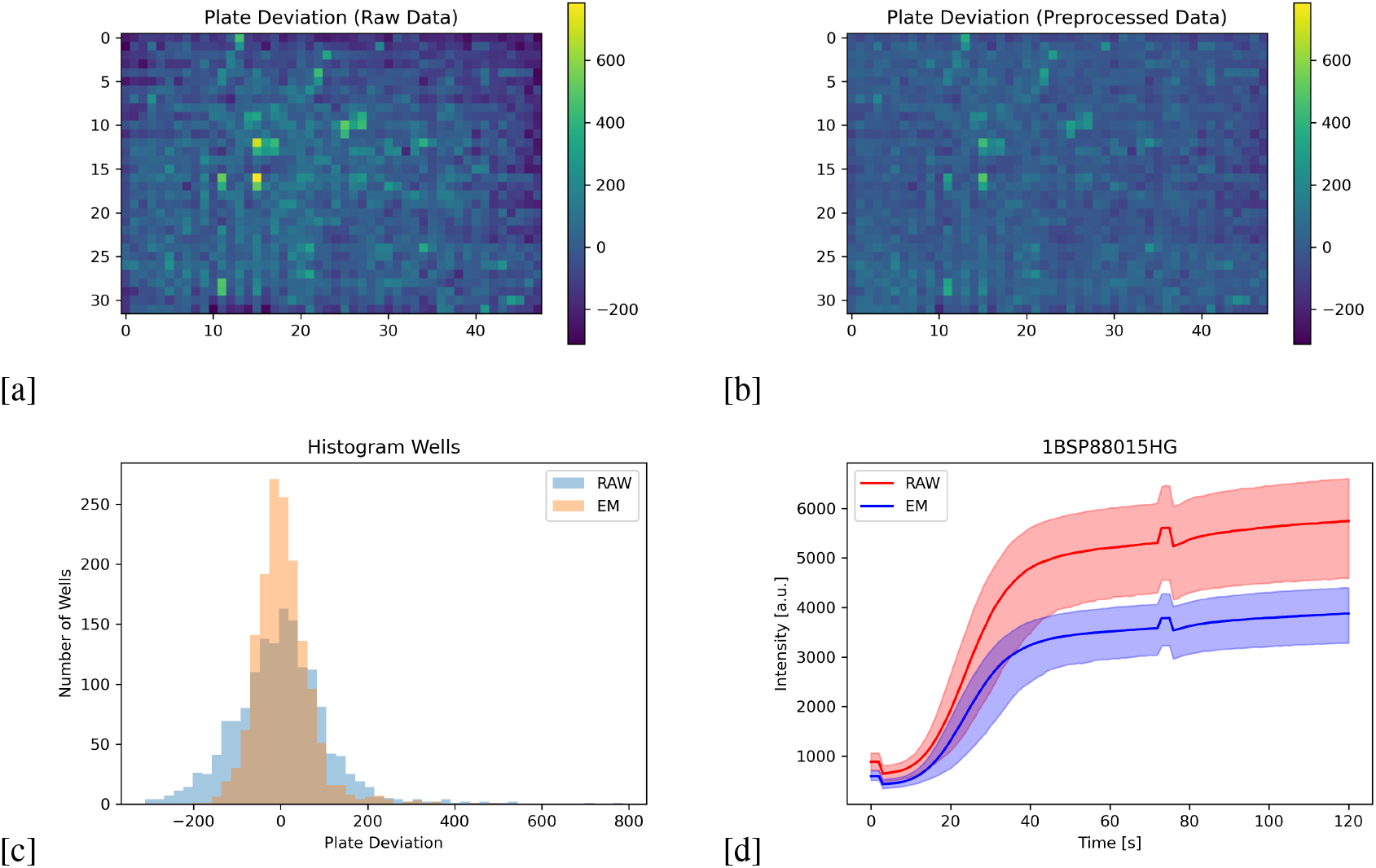
Effect of preprocessing model for plate highlighted by left arrow in Figure 3, (a) Plate-Deviation of raw signals mapped on 1536 plate; (b) Plate Deviation after preprocessing mapped on 1536 plate; (c) Distribution of Plate-Deviation for raw (blue) and preprocessed (orange) data; (d): Raw and preprocessed transient signals. Shaded area corresponds to 5th-95th percentile range.

**Figure 5.**
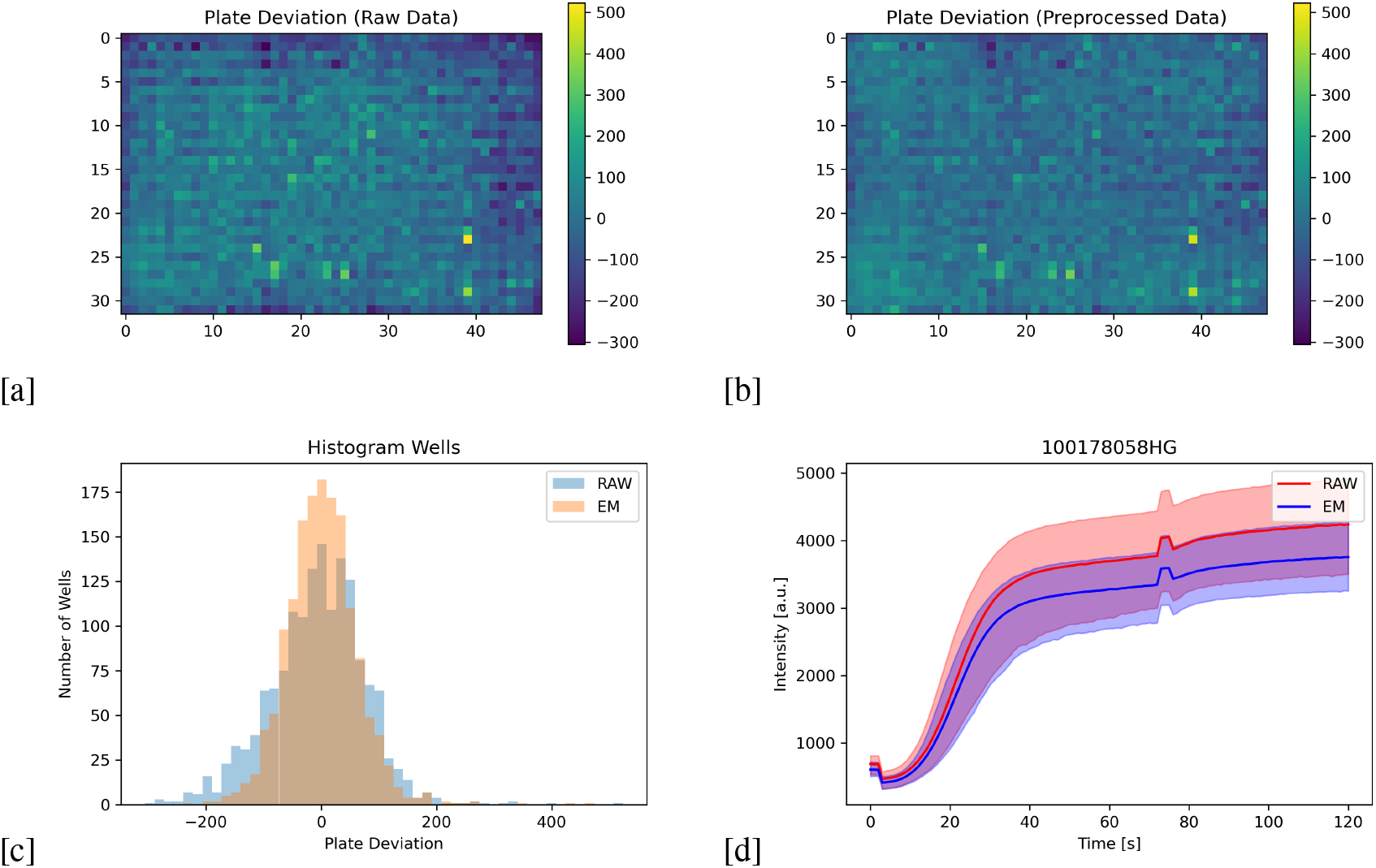
Effect of preprocessing model for plate highlighted by second to left arrow in Figure 3, (a) Plate-Deviation of raw signals mapped on 1536 plate; (b) Plate Deviation after preprocessing mapped on 1536 plate; (c) Distribution of Plate-Deviation for raw (blue) and preprocessed (orange) data; (d): Raw and preprocessed transient signals. Shaded area corresponds to 5th-95th percentile range.

**Figure 6.**
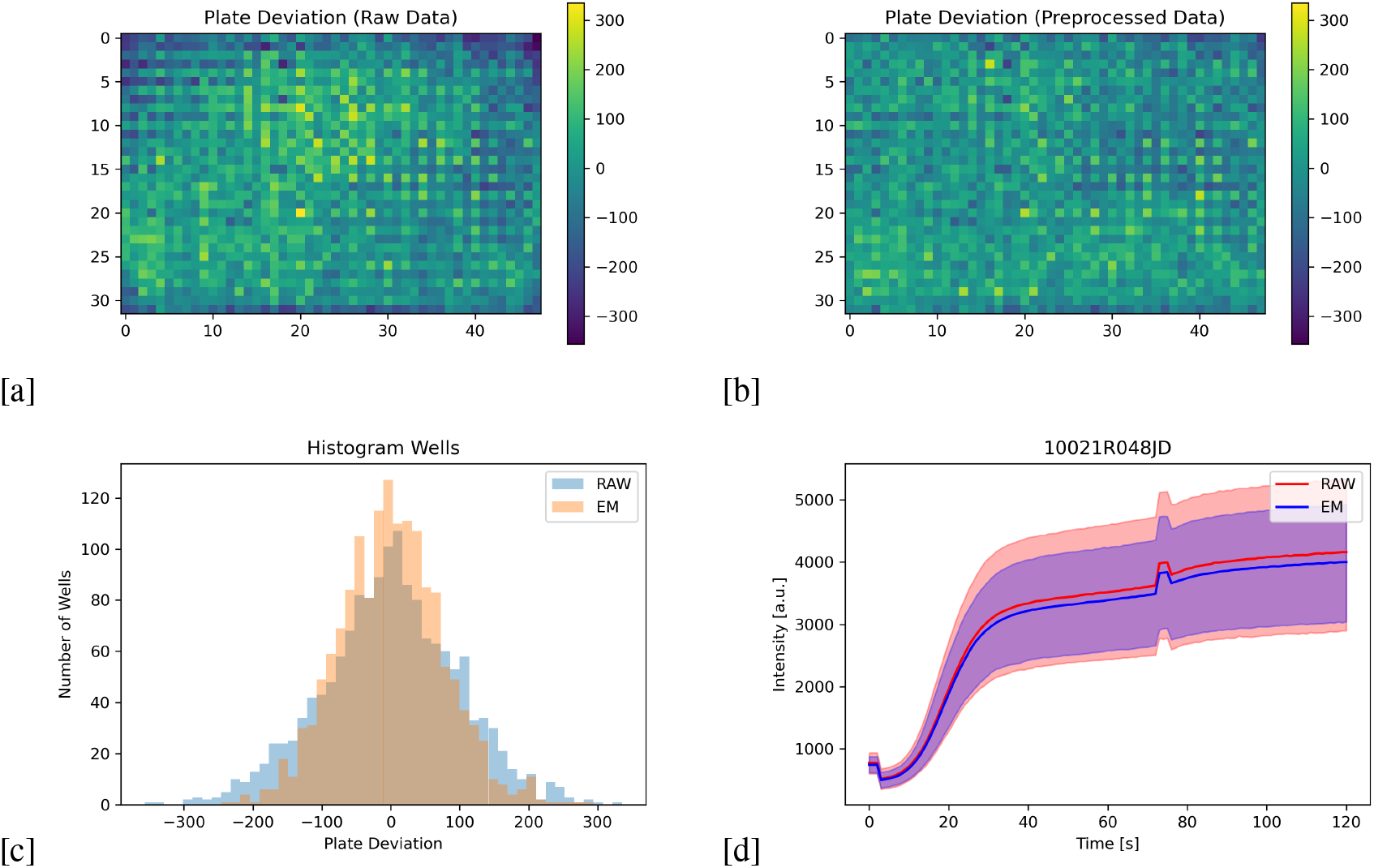
Effect of preprocessing model for plate highlighted by right arrow in Figure 3, (a) Plate-Deviation of raw signals mapped on 1536 plate; (b) Plate Deviation after preprocessing mapped on 1536 plate; (c) Distribution of Plate-Deviation for raw (blue) and preprocessed (orange) data; (d): Raw and preprocessed transient signals. Shaded area corresponds to 5th-95th percentile range.

### Dimensionality Reduction and Scoring

Figure 7 displays the two-dimensional encoding based on our analysis approach for the full uHTS data set. On this scale recognition of the global distribution of data is obstructed by the presence of strong outliers which spread out towards the second quadrant (negative latent x and positive latent y) of the two-dimensional representation. In total 112 data points reach out beyond latent x < −8 and the distribution of corresponding transient signals for these measurements is shown in Figure 7. Theses outliers exhibit a strong fluorescence response and are most likely caused by autofluorescence of the tested substances.

**Figure 7.**
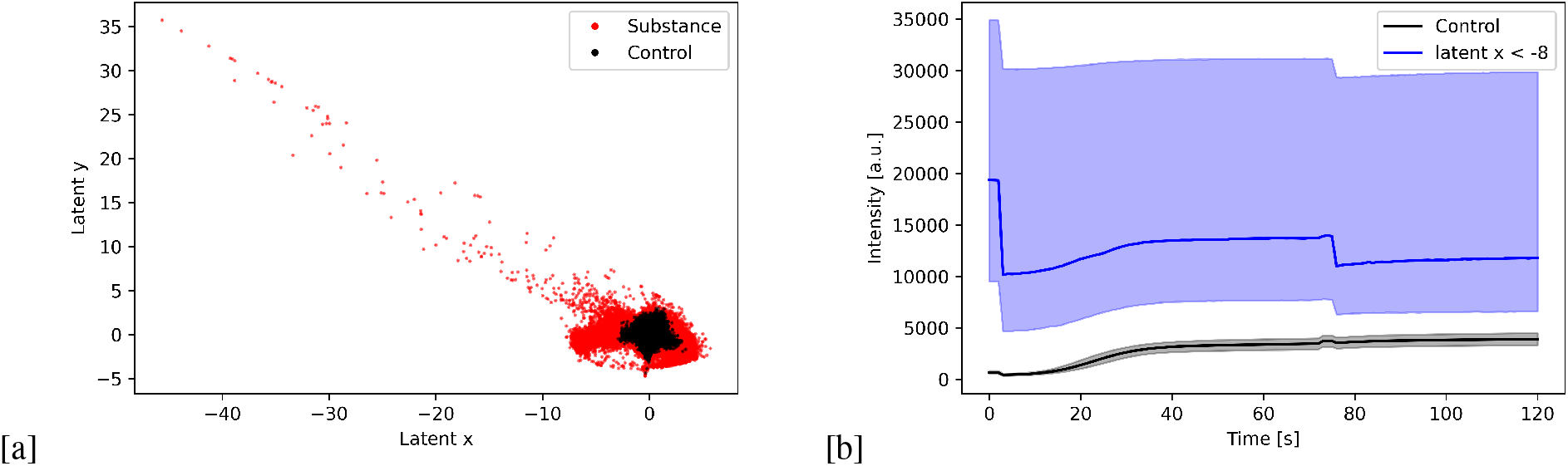
(a) Two-dimensional encoding of uHTS data set; (b) Raw signals for latent_*x*_ *< −*8

To get a clearer picture of the global distribution of data points Figure 8 highlights a region around the origin of the encoding space. Table 2 lists the 5th and 95th percentile thresholds for the distribution of points along the latent x and y axis for control and substance measurements. These percentile values clearly indicate that a vast majority of data points is located close to the origin of the two-dimensional embedding. This finding is in agreement with common experience in uHTS experiments. Primary hits are rare events and as a consequence most substances behave like control samples. A comparison of percentile thresholds for control and substance measurements further shows that control measurements are more densely packed around the origin. This fact is in line with the additional term in the loss function of our variational autoencoder model, which constrains the length of encoding vectors for known control measurements.

**Figure 8.**
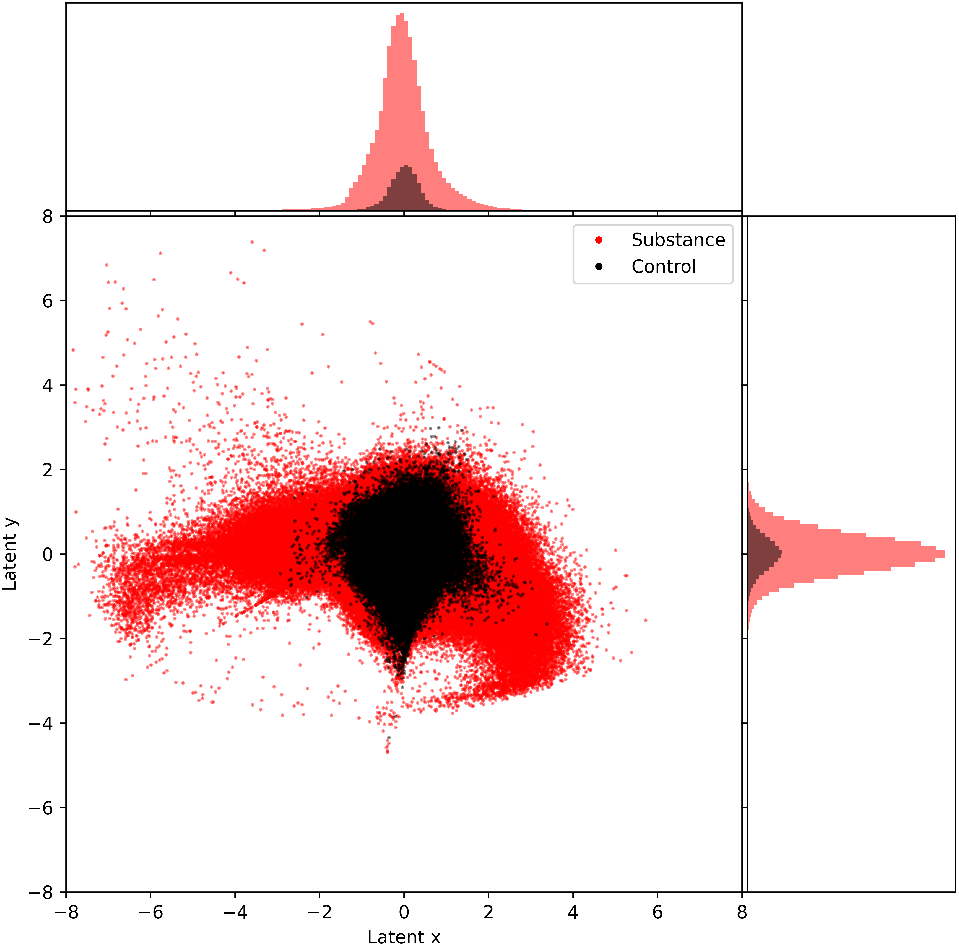
Two-dimensional encoding of uHTS data set after removal of outliers exhibiting strong fluorescence signals and distribution of data across encoding dimensions

**Table 2.** Percentile thresholds for the distribution of points along latent x and latent y axis for control and substance measurements.

| Data | 5th-percentile | 95th-percentile |
| --- | --- | --- |
| Latent x (Control) | -0.6207 | 0.5748 |
| Latent y (Control) | -0.7125 | 0.7010 |
| Latent x (Substance) | -1.0851 | 1.1682 |
| Latent y (Substance) | -0.7791 | 0.8043 |

Using contrast scores allows further dissection of the encoded data set (see supplementary material for a detailed explanation on the calculation of contrast scores). Based on calculated contrast scores for each observation we defined integer-spaced intervals and binned observations accordingly to obtain contrast categories. The number of substance samples belonging to each of the resulting categories is listed in Table 3. By removing substances belonging to the lowest contrast category (control-like observations) finer details of the data distribution can be uncovered. Six separate subfigures in Figure 9 show the distribution of substance measurements for individual contrast categories. Combining spatial information and contrast scores reveals two distinct subtypes of signals in the data set as highlighted in Figure 10. In the context of the given experiment these subtypes of signals correspond to two different modes of action of the tested substances. Reaching out along the x-axis towards negative infinity signals exhibiting a distinctively higher basal signal level are located. This response can be identified as an agonistic effect on the biological target (MoA1). In contrast to this, signals located towards the positive x-axis clearly indicate an inhibitory effect of the tested substances on the biological target (MoA2). These findings show that a combination of spatial information in the encoding space and contrast scores can be used to identify useful regions of interest in the encoding space.

**Table 3.** Distribution of contrast categories.

| Contrast Score Interval | Contrast Category | Number of Substance Samples |
| --- | --- | --- |
| < 1 | 0 | 2946558 |
| [1,2[ | 1 | 843638 |
| [2,3[ | 2 | 343763 |
| [3,4[ | 3 | 138423 |
| [4,5[ | 4 | 67184 |
| [5,6[ | 5 | 34463 |
| [6,7[ | 6 | 15021 |
| > 7 | 7 | 6982 |

**Figure 9.**
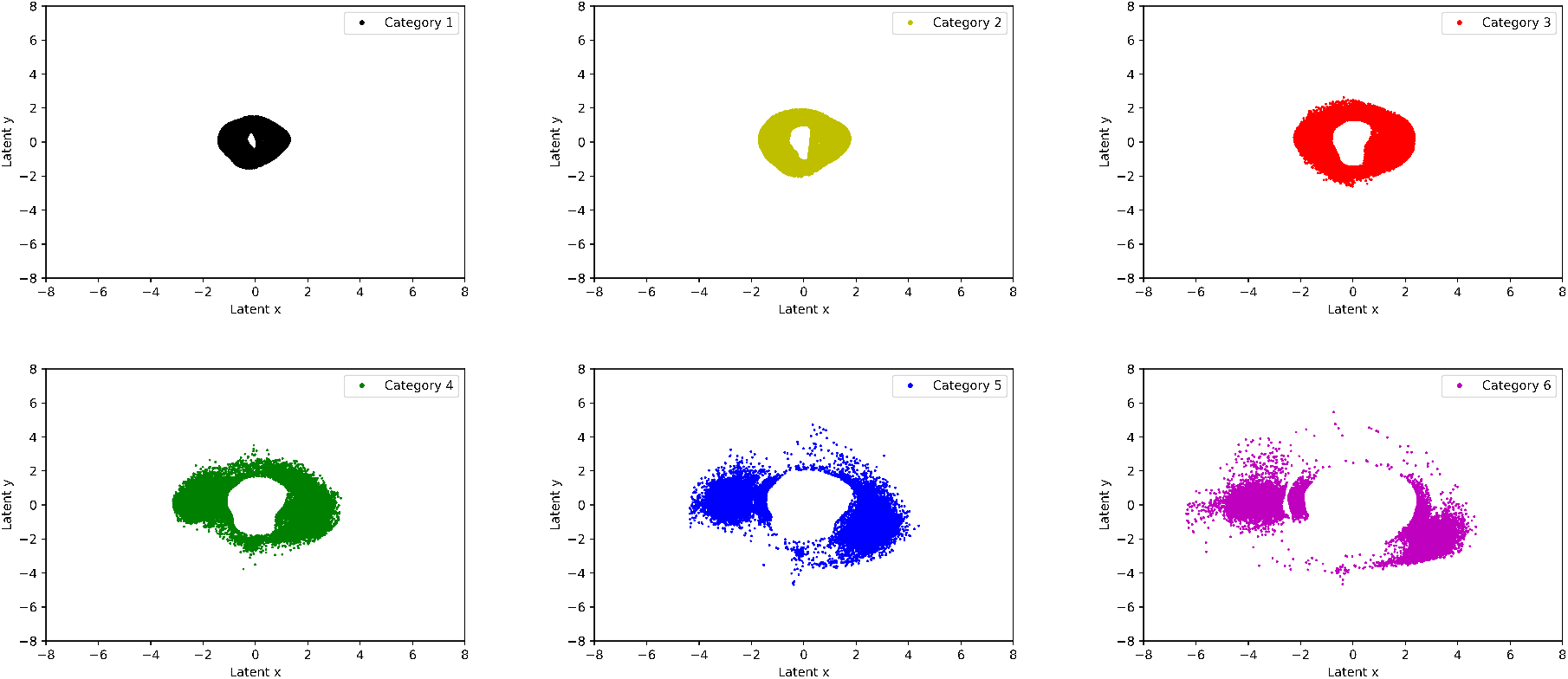
Distribution of substance measurements in embedding space for individual contrast categories.

**Figure 10.**
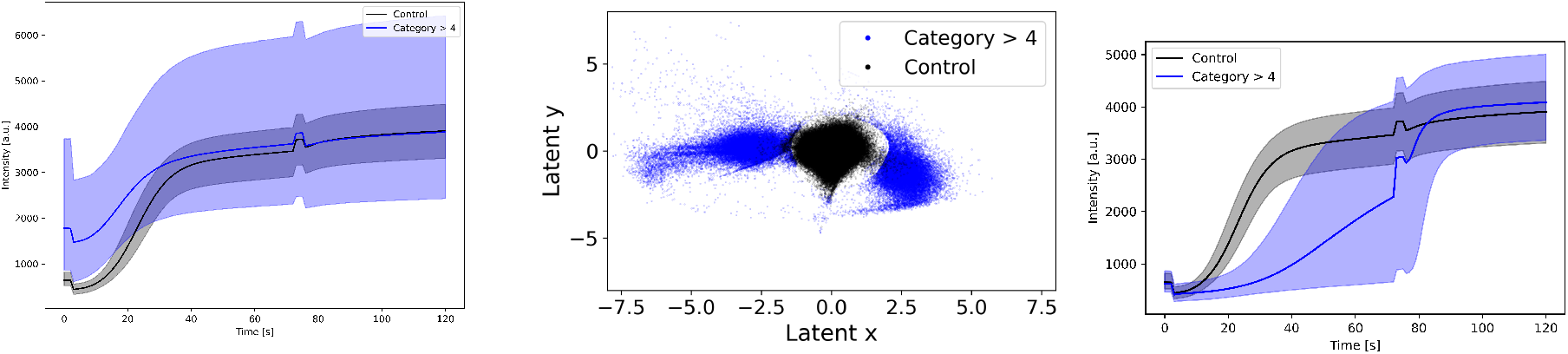
Distribution of substance measurements in embedding space for individual contrast categories.

To demonstrate the validity of the selected regions of interest we used our analysis pipeline for primary hit selection. In total 11598 substances of MoA1 and 1672 substances of MoA2 were selected and retested in a secondary screening experiment. The resulting data was processed in the same way as the primary data set and Figure 11 depicts a comparison of the encoding spaces for both data sets. In both plots MoA1 (red) and MoA2 (green) refer to the classification of compounds based on the primary screening data set. As expected, the overall distribution of data points is highly similar for both data sets and a vast majority of data points exhibit a similar response in terms of MoA classification. Definition of reconfirmation is done based on the overall distribution of contrast scores for control measurements in the secondary screening data set. As depicted in Figure 12 the 99th percentile of contrast scores for control measurements is located at 2.66. Applying this threshold to substances measurements in the data set 1501 out of 11598 MoA1 substances and 545 out of 1672 MoA2 substances are reconfirmed by the secondary screen. Figure 13 shows the distribution of these reconfirmed substances in the encoding space for the secondary screen. Interestingly, some substances apparently exhibit a different MoA compared to the primary screen classification.

**Figure 11.**
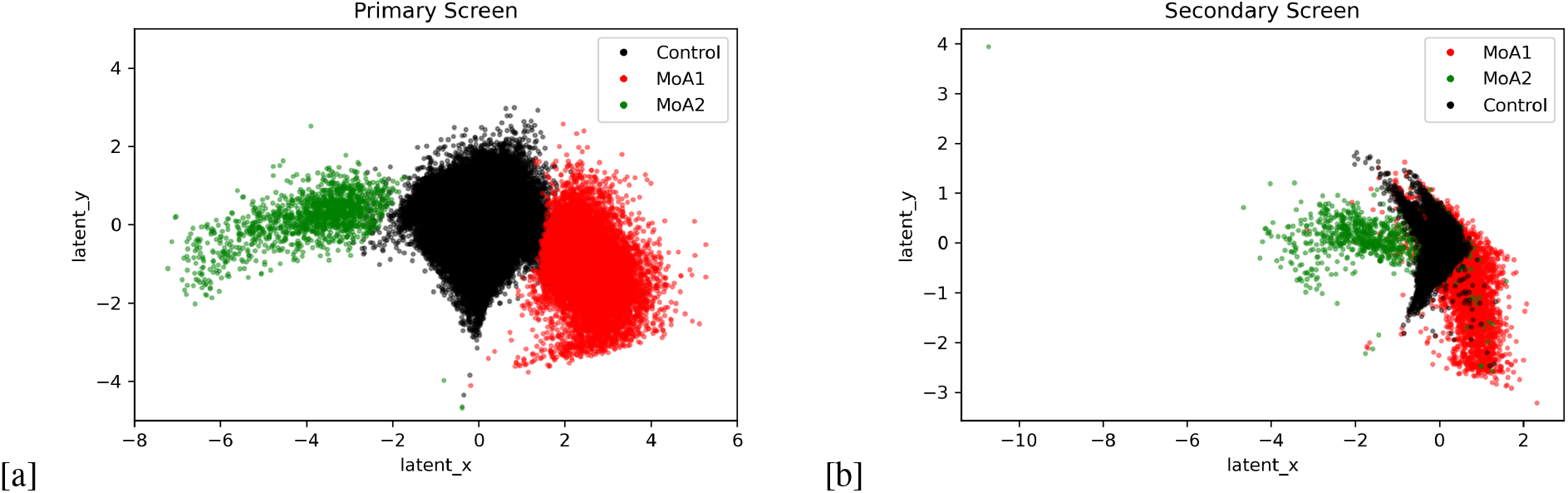
(a) MoA1 and MoA2 selected observation in encoding space of primary screening data set; (b) Encoding space of secondary screening data set for MoA1 and MoA2 compounds as defined in primary screen.

**Figure 12.**
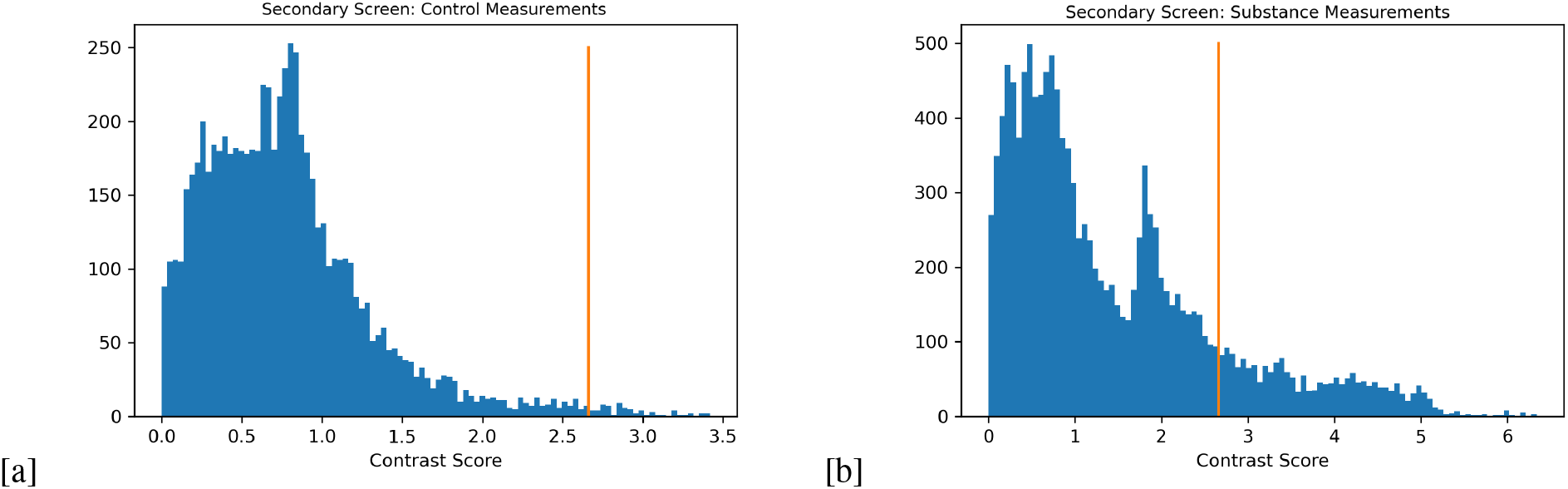
(a) Distribution of contrast scores of all control samples in secondary screen data set (orange bar: 99th quantile threshold = 2.66); (b) Distribution of contrast scores of all substance samples in secondary screen data set.

**Figure 13.**
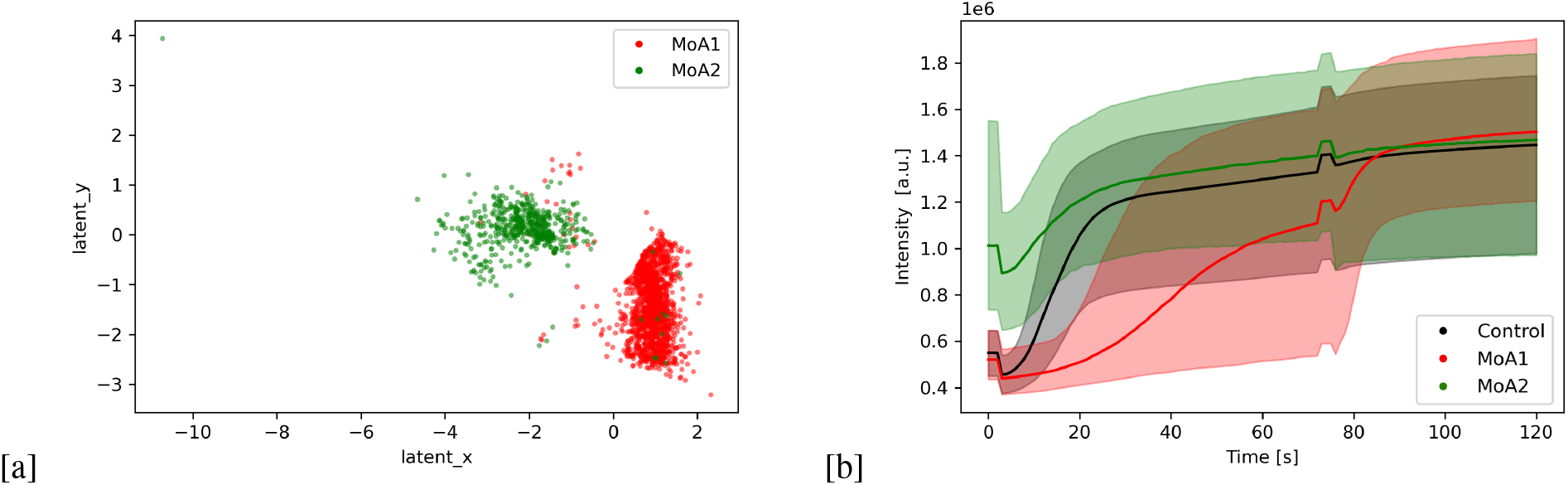
(a) Substances for MoA1 (red) and MoA2 (green) in secondary screen encoding space exhibiting a contrast score above 99th percentile of control distribution; (b) Corresponding transient signals for both MoAs and control.

### Analyzing more complex transient signals

The main strength of our approach is to derive a lower dimensional encoding of uHTS data sets without a-priori knowledge of the involved data generating mechanisms. This enables analysis of potentially much more complicated assays in which transient signals cannot easily be analyzed by point-in-time or AUC calculations. To demonstrate this, we generated a toy data set consisting of predefined subgroups of five transient signals and applied our data analysis pipeline to it. Figure 14 depicts median kinetics of all types of signals present in the toy data set. The distribution of signal classes is given in Table 4.

**Table 4.** Distribution of signal classes in generated toy data set.

| Type | Frequency of occurrences | Relative Frequency (%) |
| --- | --- | --- |
| Control | 1471734 | 95.816 |
| 1 | 12944 | 0.843 |
| 2 | 12767 | 0.831 |
| 3 | 12801 | 0.833 |
| 4 | 12829 | 0.835 |
| 5 | 12925 | 0.841 |

**Figure 14.**
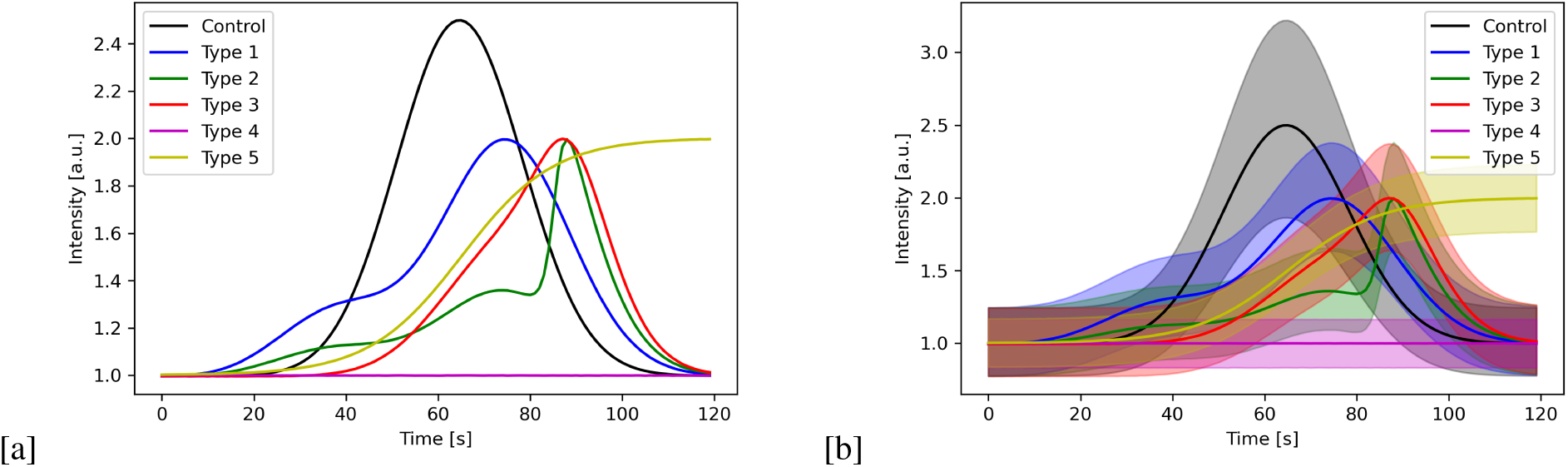
(a) Median kinetics for each signal class present in the generated toy data set; (b) 5th −95th percentile range for each signal class.

Figure 15 displays the resulting two-dimensional encoding of the data set produced by our variational autoencoder model. The results show a clear separation of all distinct types of signals present in the data set. Due to the additional term in the loss function of the autoencoder model control samples are centered around the origin of the encoding space. Further, the hierarchy of similarity between individual signal types is also captured by the generated representation, e.g. the largest distance is found for control and type 5 clusters which is in line with these signals belonging to completely different mathematical functions (gaussian type signal vs. sigmoid). On the other hand, the smallest distance is observed for type 2 and type 3 clusters.

**Figure 15.**
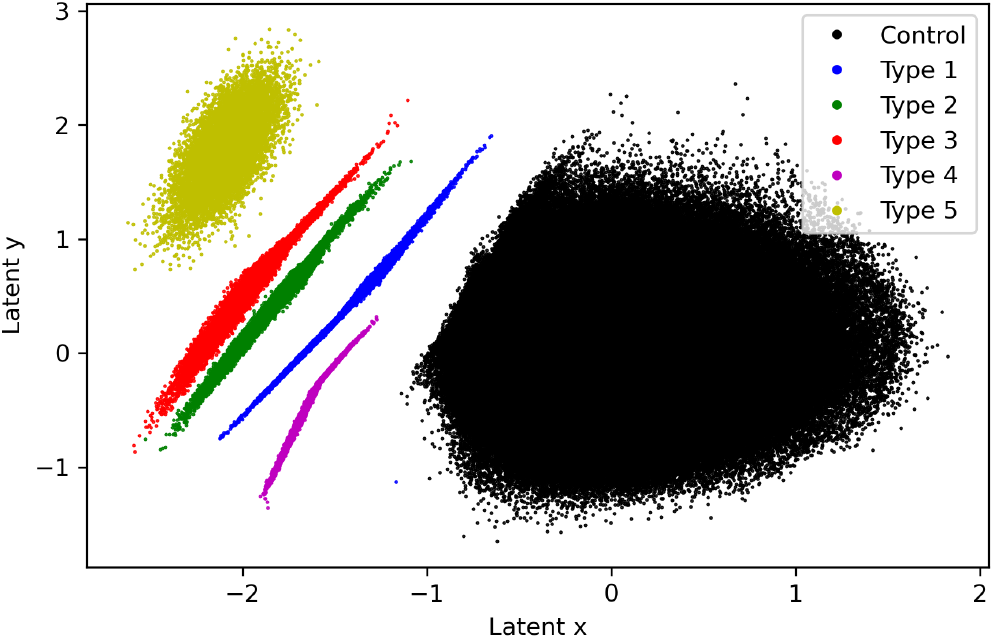
Two-dimensional encoding of toy data set.

These results also suggest a possible extension to our described analysis approach: if additional reference substances for a biological target are available, these can be used to establish alternative reference points in the encoding space. This would also enable scoring with respect to individual reference compounds in analogy to calculation of contrast scores for neutral control samples, thereby potentially allowing an accurate discrimination between different modes of actions already at primary screening level. Thus, by simple means an additional enhancement of information for screening scientists during primary hit selection can be generated using our analysis framework.

More generally, the presented deep-learning based analysis framework in this article can be viewed as a flexible starting point for analysis of uHTS data sets. Both the preprocessing model and the dimensionality reduction via the variational autoencoder can be adapted to suit experimental circumstances.

## Conclusion

In this work, we propose a novel deep-learning based analysis framework to analyze uHTS time-series data sets. The framework consists of two independent deep-learning models. A preprocessing model is used to reduce temporal and spatial signal variation caused by systematic and random errors and a variational autoencoder model is used to perform dimensionality reduction. In contrast to classical evaluation methods our approach is capable to derive lower dimensional encodings of time-series signals without a-priori knowledge of the data generating mechanism, thereby circumventing limitations of traditional point-in-time or AUC quantification methods. Including an additional term in the loss function of our variational autoencoder model the derived lower-dimensional representation of the uHTS data set contains a known point of reference. This imposed structure of the lower dimensional encoding space is leveraged to provide contrast scores for each substance sample reflecting its dissimilarity with respect to the distribution of neutral control samples. We tested our analysis framework on an experimental data set based on a cell-based calcium flux assay which has been optimized to screen for inhibitors of a biological target. Our framework indicated two distinct classes of substances in the screened library which could be attributed to two biological modes of action. Primary hits of substances belonging to both modes of action were successfully validated in a secondary screening experiment. Finally, based on a randomly generated toy data set, we illustrated the possibility to use our deep-learning based approach to evaluate even more complex uHTS data sets.

## Acknowledgements

This work was supported by Life Science Collaboration Program (Project HIDDEN) of Bayer AG.

## Author contributions statement

P.B. developed and implemented methods described in this article, wrote the manuscript and prepared all figures and tables, L.H, H.D. E.M. T.M. and D.G. ideated and designed the project, L.H. and H.D. developed prototypes for preprocessing and dimensionality reduction and guided development of methods by P.B., F.L. supported parts of the analysis and helped in the design of scoring metrics. All authors reviewed the manuscript.

## Additional information

P.B., D.G. and T.M. are employees Bayer AG, Pharmaceuticals. L.H., E.M. and H.D. are employees of Bayer AG, Crop Science. F.L. was an employee of Bayer AG, Crop Science, during all performed work related to this article. He is currently an employee of IQVIA Commercial GmbH Co. OHG.

## Data availability

The underlying proprietary screening data sets are part of Bayer’s data warehouses, which guarantees industrial and economic relevance. However, this precludes the publication of the underlying screening data sets. Screening data sets are located in controlled access data storage at Bayer AG. The artificial toy data set is available at https://doi.org/10.5281/zenodo.13605010.

