## Supplementary Material for "A deep-learning based analysis framework for ultra-high throughput screening time-series data"

### General

The pipeline is programmed in python 3.7.4 employing the standard data science software stack (numpy 1.17.2, pandas 1.1.2, scikit-learn 0.24.2[1]). All neural network models have been constructed using tensorflow 2.6.0[2].

### Dimensionality reduction by variational autoencoder

As explained in the main article the dimensionality reduction step of our analysis paves the way for defining a scoring system based on the similarity of an individual measurement with respect to control measurements. Therefore, we intend to shape the encoding space to contain a known point of reference, which is given by known control measurements within the data set. To achieve this, we added an additional term to the loss function of the variational autoencoder. Typically the loss function of a standard variational autoencoder contains two parts[3]:

- Reconstruction loss: This term tracks the quality of the reconstructed input signals in our case quantified by the mean squared error between input- and reconstruction-vectors:

$$L_{reconstruction} = \frac{1}{d}\sum_{i=1}^{d}(x_i - y_1)^2 \tag{1}$$

  where $x_i$ and $y_i$ are the $i^{th}$ component of the input- and reconstruction vectors each having $d$ components.

- Latent loss: This term quantifies the difference between the distribution of the encoding vector-components (generated by the encoder part of the model) and a given prior distribution (here we use a Gaussian prior with zero mean and unit variance). This difference is calculated via the Kullback-Leibler divergence for the two distributions. Under these circumstances the latent loss function can be expressed as[3]:

$$L_{latent} = -\frac{1}{2}\sum_{i=1}^{k} 1 + log(\sigma_i^2) - \sigma_i^2 - \mu_i^2 \tag{2}$$

  where $\mu_i$ and $\sigma_i$ are the mean and standard deviation of the $i^{th}$ component of the encoding vector which has $k$ components (in our case $k = 2$).

For our variational autoencoder model we added a third term to the standard loss function that acts to minimize the mean $\mu$ of the encoding vectors for **control samples only**. It can be expressed as:

$$L_{control} = \frac{\gamma}{k}\sum_{i=1}^{k}\mu_i^2 \tag{3}$$

where $\mu_i$ is the mean of the $i^{th}$ component of the encoding vector, $k$ gives the number of components of the encoding vector and the variable $\gamma$ takes values of 0 or 1. The value of $\gamma$ is determined by the type of input during training: 0 in case of a training example belonging to the substance measurements and 1 in case of a training example belonging to the control measurements.

The full loss function is a weighted sum of these three terms given above. Empirically, we have chosen the following weights for analysis of all the reported data sets described in the main article:

### Scoring

As mentioned in the main article, we employ an anomaly detection approach to quantify the similarity of substance measurements with respect to control measurements. Based on the two-dimensional encoding of the data set the following procedure is used to calculate contrast scores for each batch of the screening data set:

| Loss term | Weight |
|---|---|
| Latent | 0.02 |
| Reconstruction | 0.68 |
| Control | 0.3 |

**Table 1.** Weight factors for individual terms of the full loss function.

- Step 1: Density estimation of known control measurements via Gaussian-Mixtures (GMM, covariancetype $= full$) and selection of best model based on Bayesian information criterion.
- Step 2: Compute the weighted log probabilities for each sample in the batch based on the best GMM found in step 1.
- Step 3: The resulting distribution is highly skewed due to most substances being inactive. Thus, we apply a log-transformation and denote the resulting values as log-scores.
- Step 4: Calculation of contrast scores for substance measurements based on log-scores.

Detailed description of each step:

- Step 1: In order to establish a robust similarity metric we restrict our density estimation to control measurement which exhibit a distance to the origin of the encoding space below the $90^{th}$ percentile. In total, 40 GMMs are fitted to the data with an increasing number of gaussian components ($n = 1$ to 40). The best model is selected based on the Bayesian information criterion.
- Step 2 and 3: We employ scikit-learns *scoresamples* method which returns the weighted log probabilities for each sample based on the fitted GMM. Because the GMM constitutes a probability density function its values are strictly > 0 and also can exceed 1. Thus, log probabilities based on this probability density function will contain negative values as well as values > 0. Therefore, prior to log-transformation of the data we shift the data by subtracting the minimum value. Finally, numpy's *log1p* method is used to calculate log-scores.
- Step 4: The contrast score metric shall reflect the similarity of a substance measurement with respect to the distribution of neutral control measurements. Thus, we characterize this difference based on the median and interquartile range (IQR) of the distribution of log-scores for control measurements in the following way:

$$s_{contrast} = \frac{s - Median(s_{control})}{IQR(s_{control})} \tag{4}$$

where $s$ are the log-scores as calculated in step 2, $s_{control}$ refers to the log-scores of the control population and $IQR$ refers to the interquartile range.
